# R2C2: Improving nanopore read accuracy enables the sequencing of highly-multiplexed full-length single-cell cDNA

**DOI:** 10.1101/338020

**Authors:** Roger Volden, Theron Palmer, Ashley Byrne, Charles Cole, Robert J Schmitz, Richard E Green, Christopher Vollmers

**Affiliations:** Department of Biomolecular Engineering, University of California Santa Cruz, CA 95064, USA; Department of Molecular, Cellular, Developmental Biology, University of California Santa Cruz, CA 95064, USA; UC Santa Cruz Genomics Institute, Santa Cruz, California 95064, USA; Department of Genetics, University of Georgia, Athens, GA 30602, USA

## Abstract

High-throughput short-read sequencing has revolutionized how transcriptomes are quantified and annotated. However, while Illumina short-read sequencers can be used to analyze entire transcriptomes down to the level of individual splicing events with great accuracy, they fall short of analyzing how these individual events are combined into complete RNA transcript isoforms. Because of this shortfall, long-read sequencing is required to complement short-read sequencing to analyze transcriptomes on the level of full-length RNA transcript isoforms. However, there are issues with both Pacific Biosciences (PacBio) and Oxford Nanopore Technologies (ONT) long-read sequencing technologies that prevent their widespread adoption. Briefly, PacBio sequencers produce low numbers of reads with high accuracy, while ONT sequencers produce higher numbers of reads with lower accuracy. Here we introduce and validate a new long-read ONT based sequencing method. At the same cost, our Rolling Circle Amplification <u>t</u>o <u>C</u>oncatemeric <u>C</u>onsensus (R2C2) method generates more accurate reads of full-length RNA transcript isoforms than any other available long-read sequencing method. These reads can then be used to generate isoform-level transcriptomes for both genome annotation and differential expression analysis in bulk or single cell samples.

**Significance Statement:** Subtle changes in RNA transcript isoform expression can have dramatic effects on cellular behaviors in both health and disease. As such, comprehensive and quantitative analysis of isoform-level transcriptomes would open an entirely new window into cellular diversity in fields ranging from developmental to cancer biology. The R2C2 method we are presenting here is the first method with sufficient throughput and accuracy to make the comprehensive and quantitative analysis of RNA transcript isoforms in bulk and single cell samples economically feasible.

## Introduction

Short-read RNAseq has been used for the analysis of transcriptomes for over a decade (1). The massive read output of Illumina sequencers makes it possible to quantify gene expression accurately using this approach. However, to accommodate Illumina sequencers’ short read-length, RNA or cDNA has to be fragmented during sample preparation, thereby losing long distance RNA transcript isoform information. Specialized protocols like SLR(2) or spISO-seq(3) have been used successfully to recover long-distance information but they require either specialized instrumentation or complex workflows. The SLR method assembles mostly incomplete cDNA molecules, and has limited throughput, while spISO-seq requires a 10X Genomics instrument and generates read clouds which capture long distance information, and yet cannot assemble full-length cDNA molecules.

In contrast, long-read sequencing technology has the capability to sequence entire cDNA molecules end-to-end. Currently, the PacBio Iso-Seq pipeline represents a powerful gold standard for cDNA sequencing(4) and has been used to investigate a wide range of transcriptomes(5, 6). The PacBio Sequel sequencer produces ~200k accurate circular consensus reads of full-length cDNA molecules per run.

ONT technology could present a valuable alternative for cDNA sequencing, because the ONT MinION can currently generate more than one million reads per run. We and others have shown that the ONT MinION can sequence cDNA at high throughput, but that data analysis is challenging (7, 8) due to its high error rate. Base level identification of splice junction sequence is the main challenge.

One strategy to increase the base accuracy of cDNA sequences produced by the higher-output ONT MinION sequencer is to apply the circular consensus principle applied by PacBio sequencers. By sequencing 16S amplicon molecules, the INC-seq(9) method has shown that this is possible, in principle. But, the reported throughput of a few thousand reads per-run would be insufficient for transcriptome analysis. Further, like PacBio technology, the INC-Seq method uses blunt-end ligation to circularize double-stranded DNA molecules, which does not differentiate between full-length or fragmented DNA molecules. In summary, current technology produces reads that are either too inaccurate (ONT), potentially incomplete (Illumina, PacBio, ONT, INC-seq), or too low-throughput/expensive (PacBio, SLR, INC-seq) to enable high-throughput complete cDNA sequencing.

Here we introduce the <u>R</u>olling Circle <u>t</u>o <u>C</u>oncatemeric <u>C</u>onsensus (R2C2) method which overcomes these limitations by leveraging the long read length of the ONT technology to generate consensus sequences with increased base accuracy. First, we benchmark R2C2 against the PacBio Iso-Seq gold standard for the analysis of the same synthetic transcript mixture. Second, we apply R2C2 to analyze the transcriptomes of 96 single B cells derived from a healthy adult. We show that a single run of R2C2 can generate over 400,000 reads covering full-length cDNA molecules with a median base accuracy of 94%. Using a new version of our Mandalorion pipeline, these reads can be used to identify high confidence RNA transcript isoforms present in bulk or single cell transcriptomes. Illustrating the power of this approach, we find that many of the B cells in our study express RNA transcript isoforms of the CD19 gene that lack the epitope targeted by CAR T-cell therapy (10–12).

## Results

### R2C2 improves the base accuracy of the ONT MinION

To benchmark the R2C2 method, we analyzed SIRV E2 synthetic spike-in RNA. First, we reverse transcribed and amplified the synthetic spike-in RNA using the Tn5Prime(13) protocol, which is a modification of the Smart-seq2 protocol which uses a distinct template switch oligo containing 7 nucleotide sample indexes during reverse transcription. Amplification introduces an additional 8 nucleotide index into the cDNA molecules. The amplified cDNA is then circularized using a DNA splint and the NEBuilder Hifi DNA Assembly Master Mix, a proprietary variant of Gibson Assembly. The DNA splint is designed to circularize only full-length cDNA terminating on both ends in sequences complementary to the primers used to amplify cDNA (Fig. 1). Circularized cDNA is then amplified using Phi29 and random hexamers to perform Rolling Circle Amplification (RCA). The resulting High Molecular Weight (HMW) DNA was then debranched using T7 Endonuclease and sequenced on the ONT MinION sequencer using the 1D sequencing kit (LSK108) kit and R9.5 flowcell (FLO-MIN107).

**Fig. 1:**
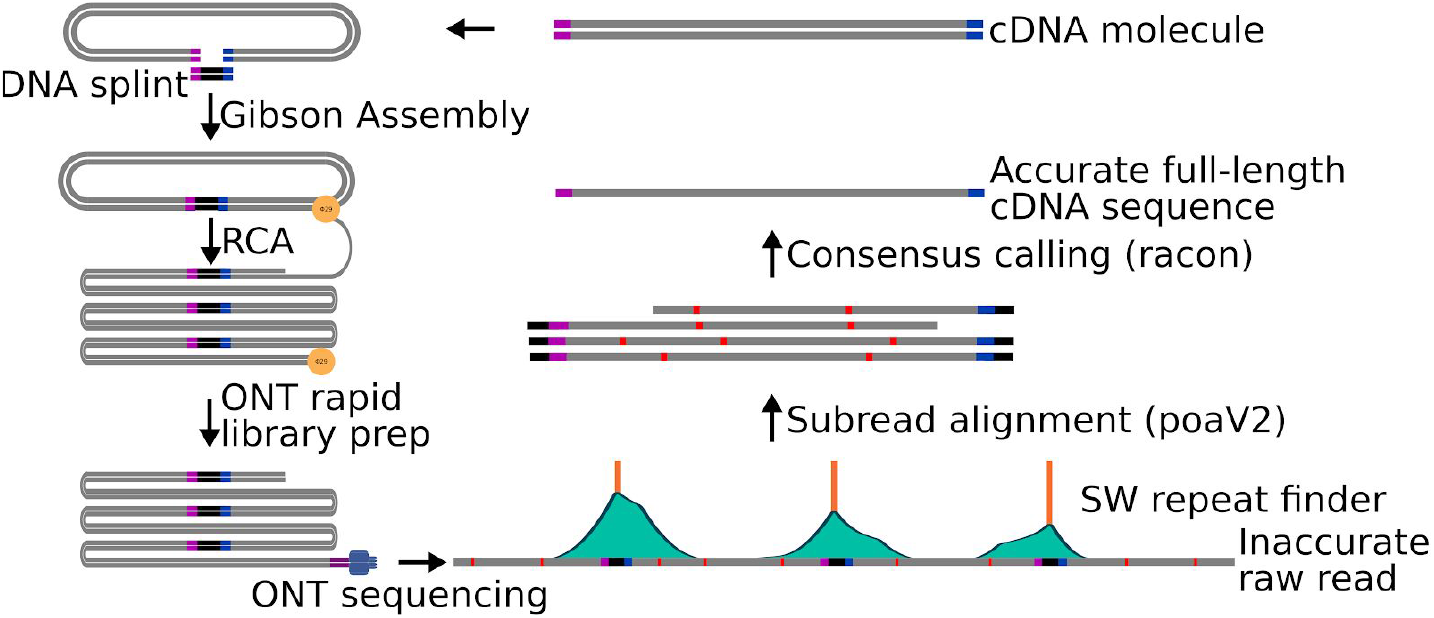
R2C2 method overview. A) cDNA is circularized using Gibson Assembly, amplified using RCA, and sequenced using the ONT MinION. The resulting raw reads are split into subreads containing full-length or partial cDNA sequences, which are combined into an accurate consensus sequences using our C3POa workflow which relies on a custom algorithm to detect DNA splints as well as poaV2 and racon.

The sequencing run produced 828,684 reads with an average length of 5.0kb resulting in a total base output of 4.1Gb. For downstream analysis we selected 621,970 of these reads that were longer than 1kb and had a raw quality score(Q) ≥ 9. We next used our C3POa (<u>C</u>ontameric <u>C</u>onsensus <u>C</u>aller using <u>POA</u>) computational workflow to generate full-length cDNA consensus reads from the raw reads. C3POa detects DNA splint sequences raw reads using BLAT(14). Because BLAT is likely to miss DNA splint sequences in the noisy raw reads, we analyze each raw read for which BLAT found at least one DNA splint sequence with a custom repeat finder which parses the score matrix of a modified Smith-Waterman self-to-self alignment (Fig 1, Fig. 2A). Repeats, or subreads, are then combined into a consensus and error-corrected using poaV2(15) and racon(16), respectively. Finally, only reads containing known priming sites at both cDNA ends are retained. In this way, C3POa generated 435,074 full-length cDNA consensus reads (and an additional 46,994 consensus reads from another multiplexed experiment) with varying subread coverage (Table 1, Fig. 2A,B).

**Fig. 2:**
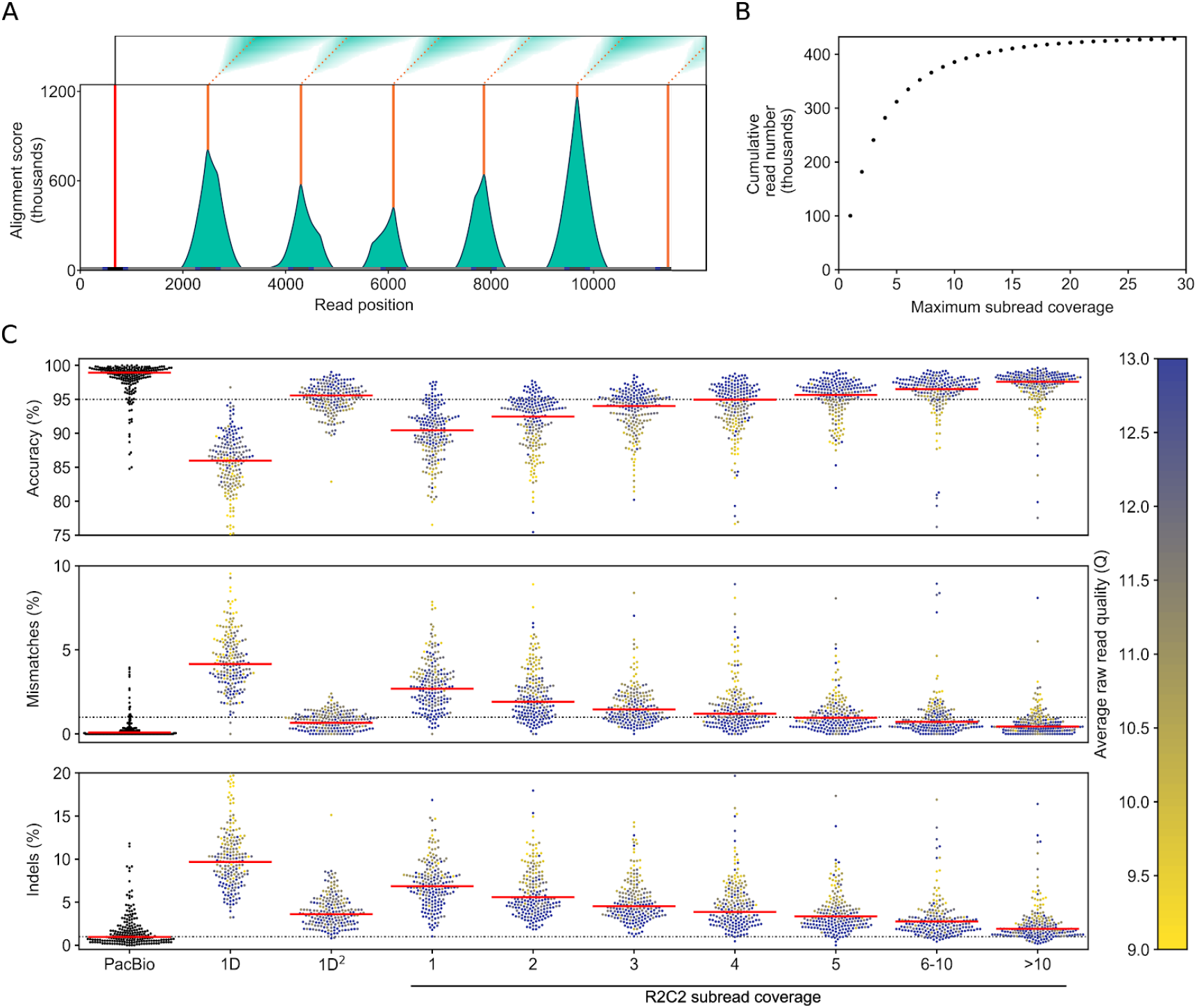
Raw reads are processed into consensus reads of varying subread coverage. A) Example of a 11.5kb raw ONT read that was analyzed by our custom Smith-Waterman repeat finder. One initial splint (red line) is identified using the BLAT aligner, then modified Smith-Waterman self-to-self alignments are performed starting from the location of the initial splint. The score matrices (on top) are then processed to generate alignment score histograms (teal). We then call peaks (orange) on these histograms. Complete subreads are then defined as the sequences between two peaks. B) Cumulative number of SIRV E2 R2C2 consensus reads is plotted against their subread coverage. C) PacBio Isoseq, standard ONT 1D, and 1D^2^ are compared to R2C2 at different subread coverage. Read accuracy is determined by minimap2 alignments to SIRV transcripts (see Methods). Median accuracy is shown as a red line. Accuracy distribution is shown as a swarm plot of 250 randomly subsampled reads. Average raw read quality of ONT reads is indicated by the color of the individual points.

**Table 1:**
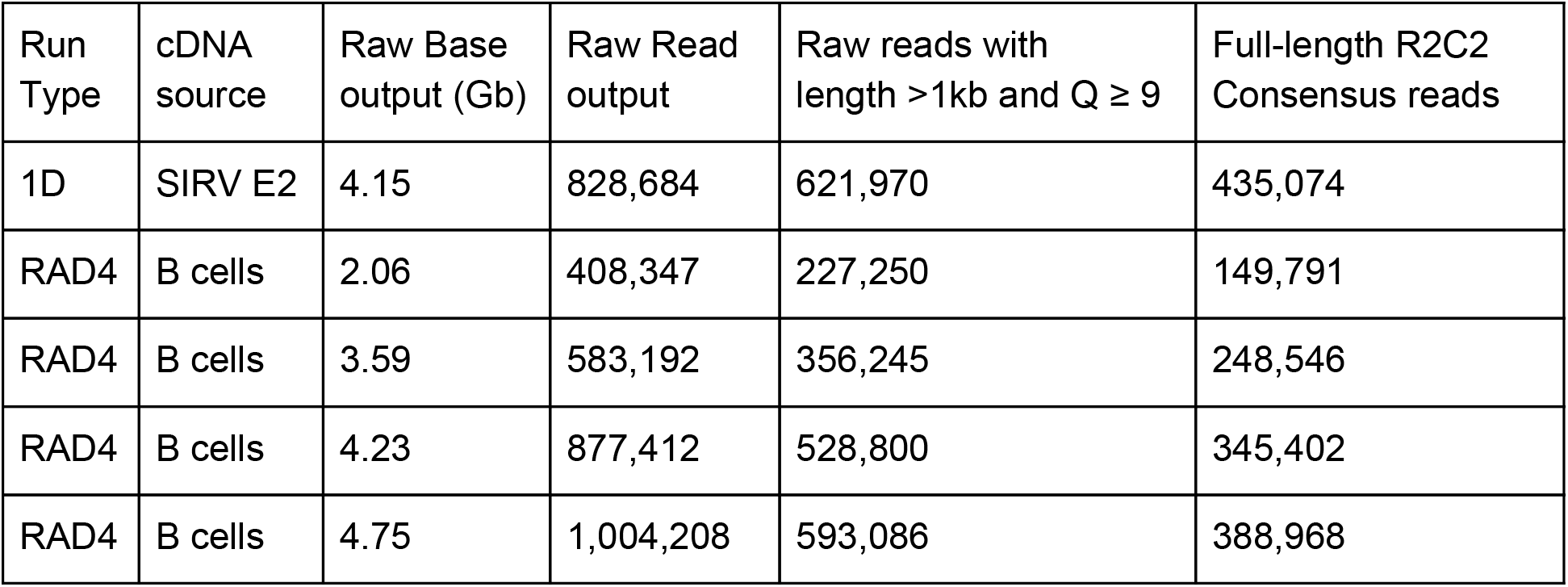
R2C2 run statistics.

We also analyzed the same cDNA pool using a standard, heavily multiplexed ONT 1D2 run generating 5,904 full-length 1D and 1,142 1D2 cDNA reads, and the PacBio IsoSeq protocol generating 233,852 full-length cDNA Circular Consensus (CCS) reads. We aligned the resulting reads generated by each protocol to the SIRV transcript sequences using minimap2 and calculated percent identity (accuracy) using those alignments. The 1D2 run produced reads with a median accuracy of 87% (1D reads) or 95.6% (1D2 reads), while PacBio CCS reads had a median accuracy of 98.9%. R2C2 reads had a median accuracy of 94% (Fig. 2C) with the accuracy of individual R2C2 reads being highly correlated with average quality score of its underlying raw read as well as the numbers of subreads this raw read contained (Fig. 2C). While mismatch errors declined rapidly with increasing number of subreads, insertion and deletion errors declined more slowly. This might be explained by insertion and deletion errors not being entirely random but systematically appearing in stretches of the same base, i.e. homopolymers (8). Indeed, 4-mers (‘AAAA’,’CCCC’,’TTTT’,’GGGG’) were enriched around insertion and deletion errors in R2C2 consensus reads (Fig. S1). Overall, more aggressive filtering of R2C2 reads based on raw read quality score and subread coverage could increase the median accuracy of the R2C2 method but would also reduce overall read output.

### R2C2 correlates well with PacBio for the quantification of SIRV transcripts

SIRV E2 transcripts vary in length from ~0.3-2.5 kb and are provided in four nominal concentration bins (“1/32”,”1/4”,”1”,”4”) varying across two orders of magnitude. By analyzing the same SIRV E2 cDNA pools using R2C2 and PacBio IsoSeq we found that our R2C2 transcript counts generally matched nominal SIRV concentrations (Fig 3A). Additionally, there seems to be no clear length bias (Fig 3B), and our R2C2 transcript counts matched PacBio transcript counts very well with a Pearson correlation coefficient of 0.93 (Fig 3C).

**Fig. 3.**
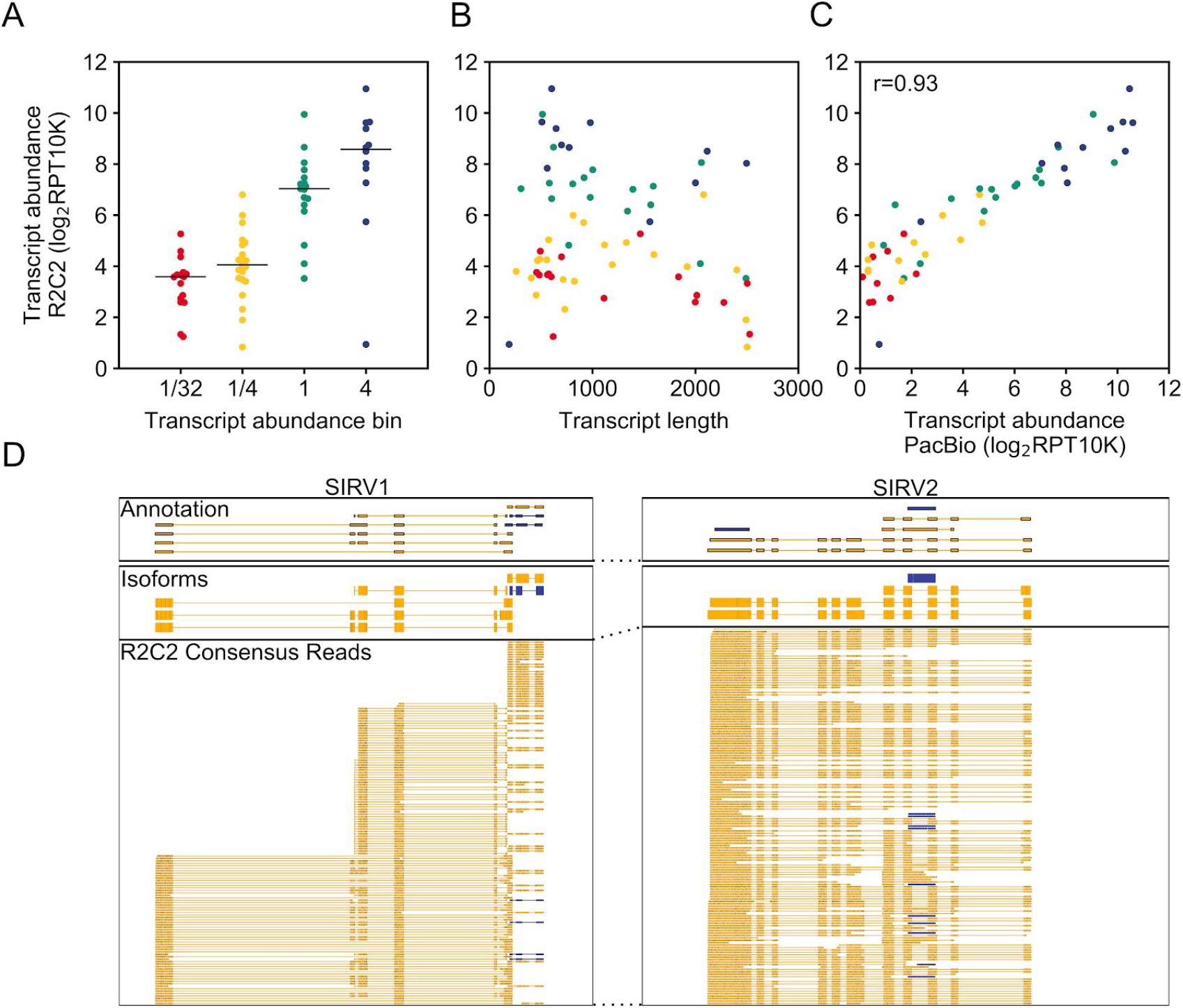
R2C2 reads can quantify SIRV transcripts. R2C2 reads were aligned to SIRV transcripts using minimap2 and expression values transcript abundance determined as Reads Per Transcript Per 10K reads (RPG10K). The transcript count ratio was plotted on the y-axis against A) the nominal transcript abundance bin reported by the SIRV transcript manufacturer (Lexogen), B) the transcript length, and C) transcript count ratio calculated from PacBio Isoseq reads. Pearson correlation coefficient (r) is reported in C). Each point represents a transcript and is colored according to it transcript abundance bin in all panels. D) Genome browser view of Transcriptome annotation, isoforms identified by Mandalorion, and R2C2 consensus reads is shown of the indicated synthetic SIRV gene loci. Transcript and read direction is shown by colors (Blue: + strand, Yellow: - strand)

This indicates that the potential variation in transcript quantification seen in Figure 3A were either rooted in differences in the initial RNA concentration found in the SIRV E2 mix or biases of our modified Smart-seq2 based cDNA amplification step rather than new biases introduced by the sequencing technology.

### R2C2 enables simple and accurate isoform identification

Next we tested whether the increased accuracy of R2C2 reads would benefit splice junction and isoform identification. To this end, we aligned PacBio, ONT and R2C2 reads to the artificial SIRVome sequence provided as a genome reference for their SIRV transcripts (Fig. 2D). 91% of splice junctions in R2C2 consensus reads matched annotated splice sites perfectly, far exceeding ONT 1D raw reads at 80% and approaching PacBio CCS reads at 96%.

This increased accuracy allowed us to simplify our Mandalorion pipeline for isoform identification (see Methods). To test how this new version of Mandalorion would perform we subsampled R2C2 consensus read alignments to levels found in highly expressed genes in whole transcriptome analysis (500 read alignments per SIRV gene locus). Some of these subsampled R2C2 consensus reads did not align from end to end to a SIRV transcript (Fig. 2D). We suspect they are products of cDNA synthesis of degraded RNA molecules likely caused by repeated freeze-thaw cycles of the SIRV E2 standards for they all contained complete 5’ and 3’ priming sites and adapter sequences. This highlighted the importance of RNA integrity for full-length transcriptome sequencing. Indeed, R2C2 reads created from single B cell lysates which are thawed only once immediately before cDNA synthesis showed evidence of degradation products at much lower levels (Fig. S2).

Because these degradation products appear to be largely random, they had little effect on the Mandalorion pipeline which identified 34 high confidence isoforms based on the subsampled R2C2 consensus reads (Fig. 2D). 24 of these isoforms matched annotated transcripts from the “1” and “4” abundance bins, while eight isoforms matched annotated transcripts from the “1/4” and “1/32” abundance bins. Only two high confidence isoforms represented truncated transcripts, caused by an oligodT mispriming on an A-rich region of the SIRV303 transcript, or a premature template switch on the (likely degraded) SIRV602 transcript, respectively. This indicated that R2C2 consensus reads paired with the Mandalorion pipeline can identify complex transcript isoforms. It also highlights the difficulty of correct identification of low abundance transcript isoforms and the abiding problem of incomplete cDNA amplification.

### R2C2 allows the demultiplexing of 7-8nt cellular indexes

Next we tested whether R2C2 reads are accurate enough to demultiplex reads based on short cellular indexes like those employed by 10X, Drop-Seq or our own Tn5Prime single cell RNAseq protocols. To this end, the SIRV cDNA we sequenced with the R2C2 method was indexed with 8 distinct combinations of a 7nt (TSO) and a 8nt (Nextera adapter) indexes. We found that we could confidently assign one 7nt and one 8nt index to 74% of R2C2 reads using a custom demultiplexing script based on Levenshtein distance between the observed sequence at the index position and our known input indexes. In 99.8% of the R2C2 assigned reads we found the combination of identified indexes matched one of the distinct combinations present in the cDNA pool.

### Analysis of 96 single B cell transcriptomes using R2C2

Having established that we could demultiplex our Tn5Prime data using R2C2 reads with very little crosstalk between samples, we sequenced cDNA from 96 single B cells which we have recently analyzed using Illumina sequencing (13). To streamline the sequencing reaction we used the ONT RAD4 (RAD004) kit which has a lower average read output than the ligation based 1D kit but has a much shorter (~20min) and, in our hands, more consistent and less error-prone workflow. Using the ONT RAD4 kit we generated 2,064,911 raw reads across 4 sequencing runs using R9.5 flowcells. C3POa generated 1,132,707 full-length R2C2 consensus reads which matched the length distribution of the sequenced cDNA closely (Figure 4A). 975,500 of the R2C2 consensus reads successfully aligned to the human genome and 730,023 of those aligned reads were assigned to single B cells based on their 7nt and 8nt cellular indexes. We found that the vast majority of those reads were complete on the 5’ end by comparing the alignment ends of these reads to transcription start sites (TSSs) previously identified (13) using Illumina sequencing. 653,410 of 730,023 (90%) reads either aligned to within 10bp of a predicted TSS (604,940 reads) or aligned within a rearranged antibody locus (48,470 reads) which makes accurate read alignment impossible.

**Fig. 4:**
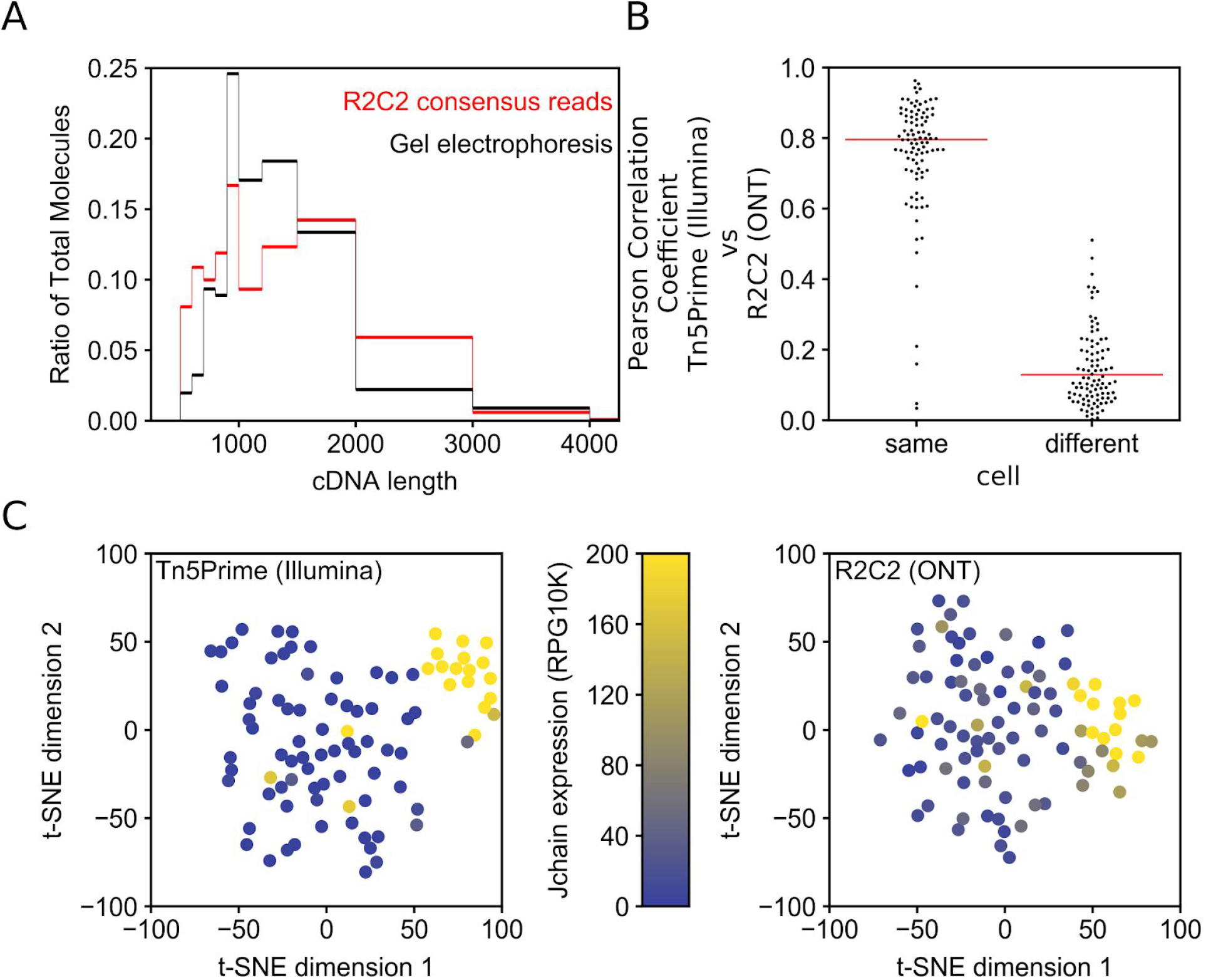
R2C2 length bias and gene expression quantification. A) B cell cDNA molecule length distribution as determined by electrophoresis on 2% agarose gel is compared to R2C2 consensus read length distribution. B) Pearson correlation coefficient (r) is shown for R2C2 and Illumina based gene expression quantification of the same of different cells. Red lines indicate medians. All 96 correlation coefficient from same cell comparisons and 96 subsampled correlation coefficients from different cell comparisons are shown as a swarmplot to display their distributions. C) t-SNE dimensional reduction plots of the same 96 B cells whose transcriptome were quantified with either the Tn5Prime Illumina based method or the R2C2 ONT based method. Cells are colored based on the Jchain expression which is strongly associated with plasmablast cell identity.

### R2C2 quantifies gene expression in single human B cells

Individual cells were assigned 7,604 reads on average. We detected an average of 532 genes per cell (at least one R2C2 consensus read overlapping with the gene). Both the numbers of genes detected as well as gene expression quantification based on these R2C2 consensus reads closely matched RNAseq-based quantification (13). When comparing gene expression of the same cell, RNAseq and R2C2 quantification had a median pearson correlation coefficient (r) of 0.79 opposed to 0.14 when comparing different cells with one another (Fig 4B). Using t-SNE clustering on R2C2 and Illumina data resulted in the sub-clustering of the same J chain-positive cells which we previously identified as plasmablasts (as opposed to memory B cells) (Fig. 4C).

### R2C2 identifies isoforms in single human B cells

We used our updated Mandalorion pipeline to identify high confidence isoforms separately for each of the 96 B cells we analyzed. By grouping R2C2 consensus reads based on their splice sites and alignment starts and ends, Mandalorion identified an average of 163 high confidence isoforms per cell. We found that identification of high confidence isoforms was dependent upon R2C2 consensus read coverage. We identified at least one isoform in 3.1%, 64.9%, 92.2% of genes covered by 1-4 reads, 5-9 reads, or >10 reads, respectively. The vast majority of genes with >10 R2C2 consensus reads contained one (78%) or two (11%) isoforms, highlighting the low complexity of single cell transcriptomes.

Overall, the isoforms we identified had a 99.1% sequence similarity with the human genome. As previously observed for mouse B1 cells (7), human B cells show a diverse array of isoforms across their surface receptors. CD37 and CD79B, which were expressed in several B cells, showed diverse isoforms. These isoforms were defined by 1) intron retention events (CD79B: Cell A12_TSO6, CD37: Cells A11_TSO2 and A17_TSO1), 2) variable transcription start sites (TSSs), and alternatively spliced exons (CD79B: Cell A20_TSO2, CD37: Cell A17_TSO1), with the alternatively spliced exon being only partially annotated (Fig.5).

Finally, for the B cell defining CD19 receptors we also observed multiple isoforms across cells, which is of particular interest because CD19 is a target for CAR T-cell therapy. Alternative splicing of CD19 has been shown to confer therapy resistance to B cell lymphomas. Interestingly, when we reference corrected (squanti-qc(17)) and translated the 4 isoforms we identified, only one contained the epitope required for FMC63 based CAR T-cell therapy (Fig. 5, Fig. S3)(10–12).

**Fig. 5.**
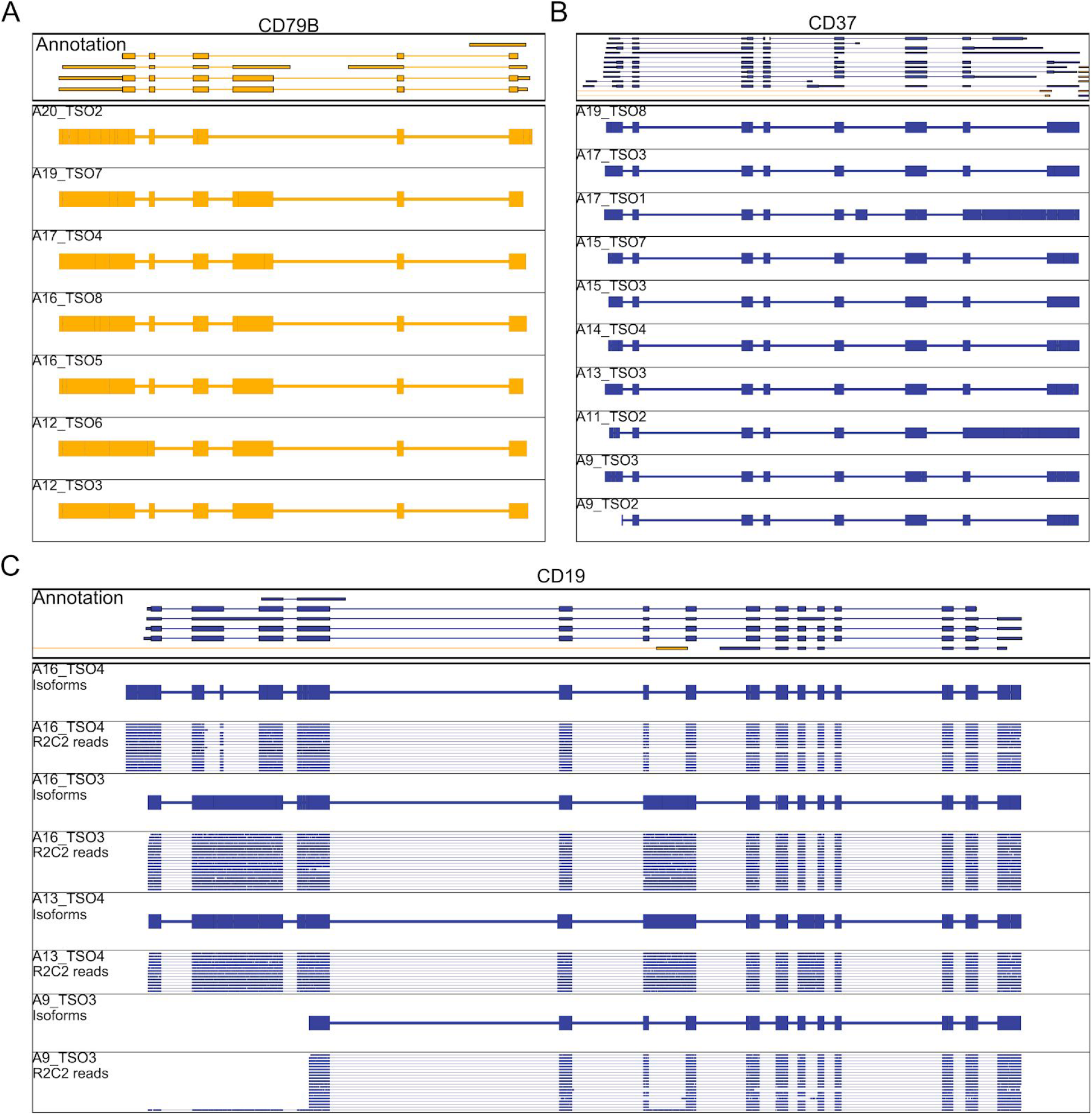
R2C2 reads identify isoforms in B cell surface receptor genes. Genome browser views of Transcriptome annotation, isoforms identified by Mandalorion, and R2C2 consensus reads (C only, downsampled to 20 reads) are shown for the indicated gene loci. Transcript and read direction is shown by colors (Blue: + strand, Yellow: - strand). Cell IDs are indicate by combinations of A and TSO indexes.

## Discussion

While RNAseq analysis has fundamentally changed how transcriptional profiling is performed, it is ultimately a stop-gap solution born from the limitations of short-read sequencing technologies. The need to fragment transcripts to fit short-read technologies like Illumina results in often unsurmountable analysis challenges. As a result, RNAseq analysis is often used like gene expression microarrays with the data used for downstream analysis being gene-expression values. Single cell RNAseq has further exacerbated this limitation because it is often restricted to 3’ or 5’ tag counting and generates gene expression values that are sparse due to both biological and technical reasons.

This results in a loss of information because individual genes can express many different isoforms, often with different functions. However, many bulk and single cell RNAseq methods do generate full-length cDNA as a intermediate product in library preparation. Long-read technology is able to take advantage of this full-length cDNA. While long-read sequencing technologies do not currently match Illumina’s read output and accuracy, their outputs and accuracies are increasing. Here, we produced over 200,000 reads at close to 99% accuracy per run using the PacBio Sequel. Further, in our hands, the standard ONT 1D^2^ protocol can generate 1 million 1D cDNA reads at 87% accuracy and 50,000 1D^2^ reads at 95% accuracy in a single run. The ONT based R2C2 sequencing method we developed takes advantage of this high throughput and increases ONT read accuracy. The R2C2 method we developed offers a compromise between PacBio and ONT technologies that generates on average 316,000 full-length cDNA reads at 94% accuracy in a single run. While the per run cost of flowcells and reagents of PacBio and ONT are roughly comparable, the capital cost of the PacBio Sequel sequencer (~$300k) vastly exceeds the cost of the ONT MinION (~$1k). This effective lack of capital costs associated with the ONT-based R2C2 method results in much lower total cost of accurate full-length transcriptome analysis compared to the PacBio IsoSeq workflow. Indeed, at its current throughput and accuracy and combined with the low cost of the ONT MinION we believe that R2C2 brings comprehensive full-length transcriptome analysis within reach of most molecular biology laboratories.

In the immediate future, the R2C2 method will be a suitable complement for short-read sequencing. To this end, the R2C2 can be easily adapted to any RNAseq library preparation protocol that produces full-length double stranded cDNA molecules with known adapter/primer sequences at their ends. This includes standard Smart-seq2, 10X Genomics, and Drop-seq protocols. Adapting R2C2 to these protocols only requires the generation of a compatible DNA splint by modifying the primers used for amplifying the DNA splint. The same cDNA pool can then be sequenced by both Illumina and R2C2 methods.

We believe that R2C2 has the potential to replace short-read RNAseq and its shotgun approach to transcriptome analysis entirely, especially considering the impending wide release of the high-throughput ONT PromethION sequencer. This will be a significant advance considering the strength of full-length transcriptome sequencing showcased here. R2C2 paired with Mandalorion accurately identified full-length synthetic transcripts as well as several surface receptor isoforms of CD79B, CD37, and CD19 expressed by 96 distinct single human B cells. Identifying these full-length isoforms with short read RNAseq would have been impossible. Finally, the CD19 RNA isoforms we identified in the single B cells derived from a healthy adult may have implications regarding immunotherapy efficacy for most lacked the epitope in exon 4 that is targeted by FMC63 based CAR T-cell therapy. This confirms that even healthy individuals contain RNA isoform diversity for CD19 which may ultimately contribute to immunotherapy resistance when undergoing FMC63 based CAR T-cell therapy (10, 11).

## Acknowledgments

We want to thank the Georgia Genomics And Bioinformatics Core (GGBC) for PacBio Sequencing Services.

## Methods

100pg of SIRV E0 (Lexogen) RNA or lysed single B cells (Collected from the blood of a fully consented healthy adult in a study approved by the Institutional Review Board (IRB) at UCSC) were amplified using the Tn5Prime(18) method, which represents a modification of the Smart-seq2(19, 20) method developed to capture 5’ ends of transcripts using Illumina sequencing.

This method uses distinct template switch oligo (TSO) and oligodT primer sequences, enabling the easy differentiation of transcript 5’ and 3’ ends when using long-read sequencing. Following the Tn5Prime protocol, RNA or Single Cell Lysate were reverse transcribed (RT) using Smartscribe Reverse Transcriptase (Clontech) in a 10ul reaction including an oligodT primer and a Nextera A TSO containing a 7 nucleotide sample index (Table S1). RT was performed for 60 min at 42°C. The resulting cDNA was treated with 1 ul of 1:10 dilutions of RNAse A (Thermo) and Lambda Exonuclease (NEB) for 30min at 37°C. The treated cDNA was then amplified using KAPA Hifi Readymix 2x (KAPA) and incubated at 95°C for 3 mins, followed by 15 cycles for SIRV RNA or 27 cycles (single B cells) of (98°C for 20 s, 67°C for 15 s, 72°C for 4 mins), with a final extension at 72°C for 5 mins. cDNA amplification requires both the ISPCR primer and a Nextera A Index primer, which contains another 8 nucleotide sample index.

### SIRV RNA

8 SIRV E2 RNA aliquots were reverse transcribed and amplified in separate reactions adding distinct 7 nucleotide TSO and 8 nucleotide Nextera A Indexes to each resulting cDNA aliquot. The separate aliquots were used directly as input into our R2C2 method or amplified using KAPA Hifi Readymix 2x (KAPA) (95°C for 3 mins, followed by 15 cycles (98°C for 20 s, 67°C for 15 s, 72°C for 4 mins), with a final extension at 72°C for 5 mins with ISPCR and Nextera_A_Universal Primers and pooled at equal amounts for input into PacBio Iso-Seq pipeline at the University of Georgia Athens sequencing core.

### Single B cell lysates

Single B cells in separate in the wells of a 96 well plate were reverse transcribed using a distinct 7 nucleotide TSO index for each row. Columns were then pooled and amplified, using a distinct 8 nucleotide Nextera A Index for each pool. This resulted in the cDNA of all 96 cells carrying a unique combination of TSO and Nextera A index. This cDNA was then pooled for Illumina sequencing (HiSeq4000 2×150)(13) or amplified using KAPA Hifi Readymix 2x (KAPA) (95°C for 3 mins, followed by 15 cycles (98°C for 20 s, 67°C for 15 s, 72°C for 4 mins), with a final extension at 72°C for 5 mins with ISPCR and Nextera_A_Universal Primers for input into our R2C2 method.

## DNA splint amplification

A ~200bp DNA splint to enable Gibson Assembly(21) circularization of cDNA was amplified from Lambda DNA using KAPA Hifi Readymix 2x (KAPA) (95°C for 3 mins, followed by 25 cycles (98°C for 20 s, 67°C for 15 s, 72°C for 30 s) using primer Lambda_F_ISPCR(RC) and Lambda_R_NextA(RC) (Table S1). This generated a double stranded DNA with matching overlaps to full-length cDNA.

### R2C2 sample preparation

#### Circularization of cDNA

200ng of cDNA was mixed with 200ng of DNA splint. Volume was adjusted to 6ul and 6ul of 2x NEBuilder Hifi DNA Assembly Master Mix (NEB) was added. The reaction was incubated for 60min at 55°C. Volume was adjusted to 20ul and non-circularized DNA was digested using 1ul of 1:10 Exonuclease III and Lambda Exonuclease as well as 1ul of Exonuclease I (all NEB). Circularized DNA was extracted using SPRI beads with a size cutoff to eliminate DNA <500bp (0.8 beads:1 sample) and eluted in 50ul of ultrapure water.

#### Rolling circle amplification

Circularized DNA was split into 5 aliquots of 10ul and each aliquot was amplified in its own 50ul reaction containing Phi29 polymerase (NEB) and exonuclease resistant random hexamers (Thermo) (5ul of 10x Phi29 Buffer, 2.5ul of 10uM(each) dNTPs, 2.5ul random hexamers (10uM), 10ul of DNA, 29ul ultrapure water, 1ul of Phi29). Reaction were incubated 30°C overnight. All reaction were pooled and volume was adjusted to 300ul with ultrapure water. DNA was extracted using SPRI beads with a size cutoff to eliminate DNA <2000bp (0.5 beads:1 sample). At this point the <u>H</u>igh <u>M</u>olecular <u>W</u>eight DNA can easily shear. Therefore, beads and samples were mixed by gentle flicking of the tube, not vortexing or vigorous pipetting. Beads were allowed to settle for 5min on magnet, and after two 70% Ethanol washes, a mix of 90ul of ultrapure water, 10ul NEB buffer 2 and 5ul T7 Endonuclease was added to the beads. Beads were incubated for 2 hour on a thermal shaker at 37°C under constant agitation. Beads were then placed on magnet and supernatant is recovered. The DNA in the supernatant is then extracted again using SPRI beads with a size cutoff to eliminate DNA <2000bp (0.5 beads:1 sample) and eluted in 15ul of ultrapure water.

1ul of the eluate was diluted in 19ul of ultrapure water. 1ul of the 1:20 dilution was used to determine the concentration of the eluate using a Qubit High Sensitivity DNA kit (Thermo). The other 19ul were analyzed on a 1% agarose gel. Successful RCA and debranching by T7 Endonuclease results in HMW DNA that runs above the 10kb band of the NEB 2-log ladder but is not stuck in the loading well.

### ONT sequencing

SIRV E2 RCA product was sequenced using the ONT 1D sample prep kit and a single 9.5 flowcell according to manufacturer’s instructions with the exception that DNA was not sheared prior to library prep. Single B cell RCA product was sequencing using the ONT RAD4 kit and four 9.5 flowcells. The resulting raw data was basecalled using the albacore (version 2.1.3) read_fast5_basecaller script with the following settings:

#### 1D run

~~~
*read_fast5_basecaller.py -r --flowcell FLO-MIN107 --kit SQK-LSK108 --output_format
fastq --input /path/to/raw_data --save_path /path/to/output_folder --worker_threads 20*
~~~

#### RAD4 runs

~~~
*read_fast5_basecaller.py -r --flowcell FLO-MIN107 --kit SQK-RAD004 --output_format fastq --input /path/to/raw_data --save_path /path/to/output_folder --worker_threads 20*
~~~

### C3POa data processing

#### Pre-processing *(C3POa_preprocessing.py)*

Basecalled raw reads underwent pre-processing to shorten read names and remove short (<1000kb) and low quality reads (Q<9) reads. Raw reads were first analyzed using BLAT(14) to detect DNA splint sequences. If one or more splint sequences were detected in a raw read, the raw read underwent consensus calling.

#### Consensus calling (*C3POa.py*)

C3POa.py is a wrapper script that performs the following steps:

1.) Tandem repeats in each raw read are detected using a modified EMBOSS WATER(22–24) Smith Waterman self-to-self alignment. First, we set the ascending diagonal of the self-to-self alignment score matrix to 0, then we sum values across the all lines parallel to the diagonal. To speed up this self-to-self alignment, the score matrix is calculated in 1000 nucleotide bins. We then call peaks along these values which indicate the position of other splint sequences in the tandem repeats the raw read contains (Fig. 1B).

2.) Raw reads are then split into complete subreads containing full repeats and incomplete subreads containing partial repeats at the read ends. If there are more than 1 complete subreads, these complete subreads are aligned using poaV2(15) with the following command:

~~~
*poa -read_fasta path/to/subreads.fasta -hb -pir path/to/alignments.pir
-do_progressive NUC.4.4.mat >./poa_messages.txt 2>&1*
~~~

The preliminary consensus is either reported by poaV2 (more than 2 subreads) or determined based on the poaV2 alignment by a custom script taking raw read quality scores into account (2 subreads). If only one complete subread is present in the raw read, its sequenced is used as consensus in the following steps.

3.) Complete and incomplete subreads are aligned to the consensus sequence using minimap2(25) and the following command

~~~
*minimap2 --secondary=no -ax map-ont path/to/consensus.fasta path/to/subreads.fastq >
path/to/subread_overlap.sam 2> ./minimap2_messages.txt*
~~~

4.) These alignments are used as input to the racon(16) algorithm which error-corrects the consensus.

~~~
*racon --sam --bq 5 -t 1 path/to/subreads.fastq path/to/subread_overlap.sam path/to/consensus.fasta path/to/corrected_consensus.fasta > ./racon_messages.txt 2>&1*
~~~

To speed up consensus calling, we divided raw reads into bins of 4000 and used GNU Parallel(26) to run multiple instances of C3POa.py.

#### Post-processing (*C3POa_postprocessing.py*)

ISPCR and Nextera Sequences are identified by BLAT and the read is trimmed to their positions and reoriented to 5’->3’.

#### Alignment

Trimmed, full-length R2C2 reads and PacBio reads are aligned to the appropriate genomes and transcripts using minimap2. The following settings were used when:

##### Aligning to SIRV transcript sequences

~~~
*minimap2 --secondary=no -ax map-ont /path/to/SIRV_Transcriptome_nopolyA.fasta
path/to/trimmed_corrected_consensus.fasta > path/to/aligned.out.sirv.sam*
~~~

##### Aligning to the “SIRVome” sequences

~~~
*minimap2 --splice-flank=no --secondary=no -ax splice /path/to/SIRVome.fasta
path/to/trimmed_corrected_consensus.fasta > path/to/aligned.out.sirvome.sam*
~~~

##### Aligning to the human genome (only chromosomes, no alternative assemblies, etc…)

~~~
*minimap2 --secondary=no -ax splice /path/to/hg38_chromosomes_only.fasta
path/to/trimmed_corrected_consensus.fasta > path/to/aligned.out.hg38.sam*
~~~

Percent identity of sequencing reads were calculated from minimap2 alignments. First md strings were added to the sam files generated by minimap using samtools calmd functionality.

Matches, mismatches and indels are then calculated based on CIGAR and md string and percent identity is reported as *(matches/(matches+mismatches+indels))*100*.

For isofom identification and visualization SAM files were converted to PSL file format using the jvarkit sam2psl (27) script.

#### Isoform identification and quantification

Isoforms were identified and quantified using a new version of the Mandalorion pipeline (EII) with the following settings:

##### Isoform Identification

~~~
*python3 defineAndQuantifyWrapper.py -c path/to/content_file -p path/to/output/ -u 5
-d 30 -s 200 -r 0.05 -R 3 -i 0 -t 0 -I 100 -T 60 -g /path/to/genome_annotation.gtf -f
/path/to/config_fi le*
~~~

~~~
*gmap -f psl -B 5 -t 6 -n 1 -d /path/to/human_reference_index
path/to/isoform_consensi.fasta > path/to/isoform_consensi.psl*
~~~

### Availability

C3POa and Mandalorion will be available at github under https://github.com/rvolden/C3POa and https://github.com/rvolden/Mandalorion-Episode-II, respectively.

Raw read data will be available upon publication at the SRA under PRJNA448331 (SIRV E2) and PRJNA415475 (B cells).

## Supplemental Figures and Tables

**Fig. S1.**
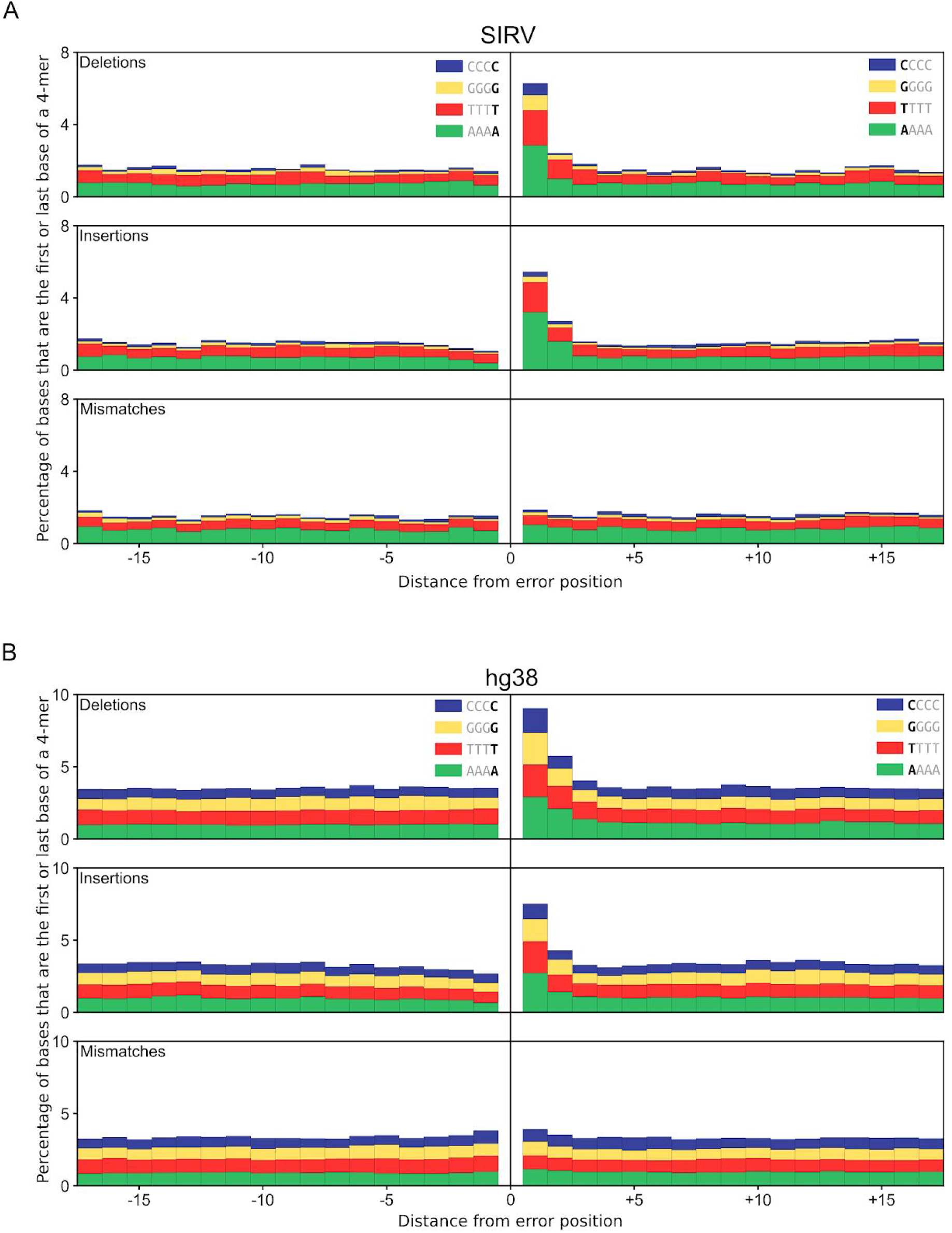
Sequence context of error in R2C2 reads. The occurrence of 4-mers (CCCC,GGGG,TTTT, or AAAA) around R2C2 sequencing deletions, insertions, and mismatches is shown as stacked bar plots for R2C2 reads covering SIRV cDNA (A) or human B cell cDNA (aligned to hg38) (B).

**Fig. S2:**
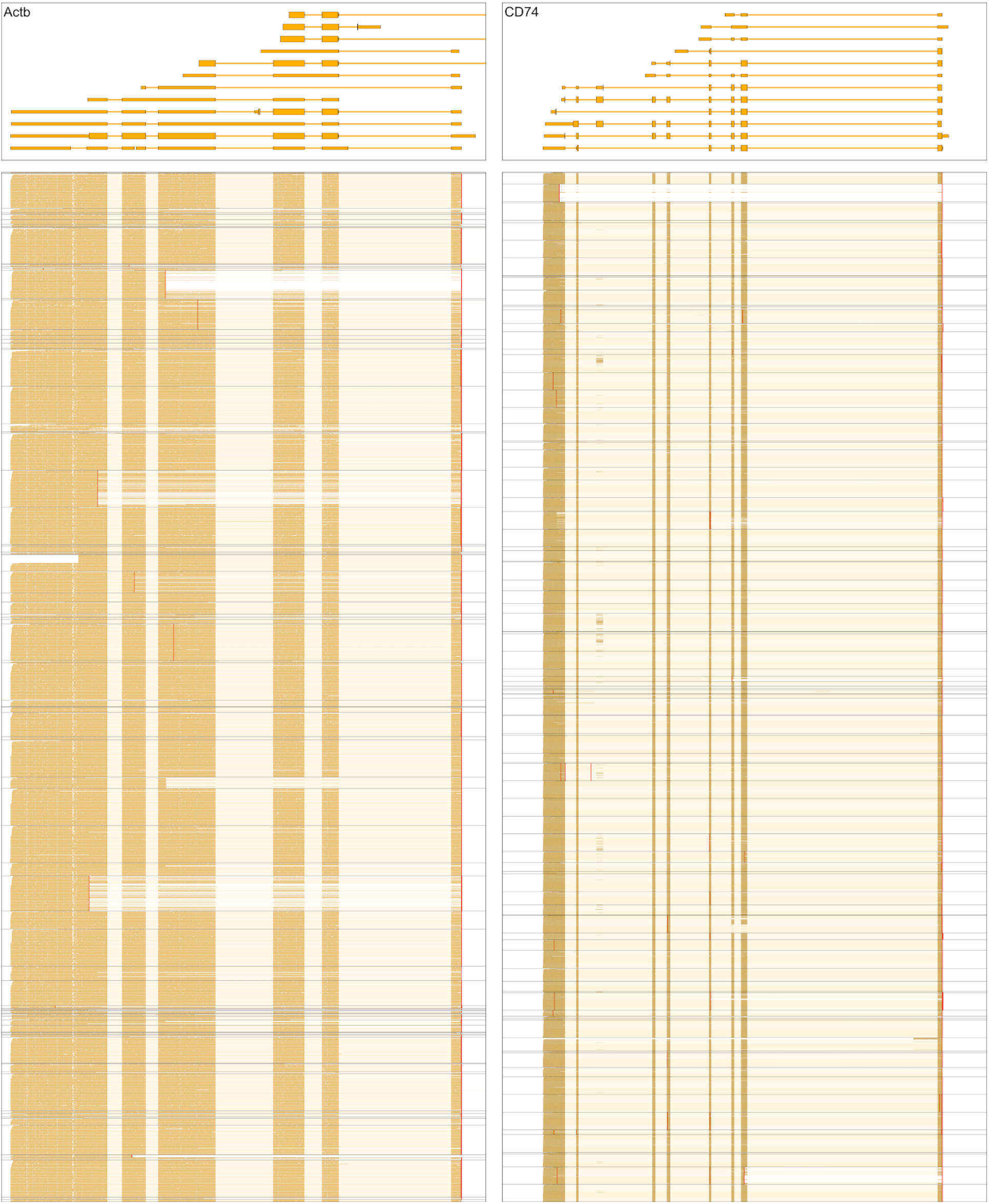
R2C2 consensus reads of single cell cDNA. Genome browser view of Transcriptome annotation and up to 200 R2C2 consensus reads per cell is shown of the indicated human gene loci. Data R2C2 consensus reads from different cell are separated by black lines. Transcript and read direction is shown by colors (Blue: + strand, Yellow: - strand). Red lines indicated TSSs identified based on the same cDNA using Tn5Prime based Illumina sequencing.

**Fig. S3:**
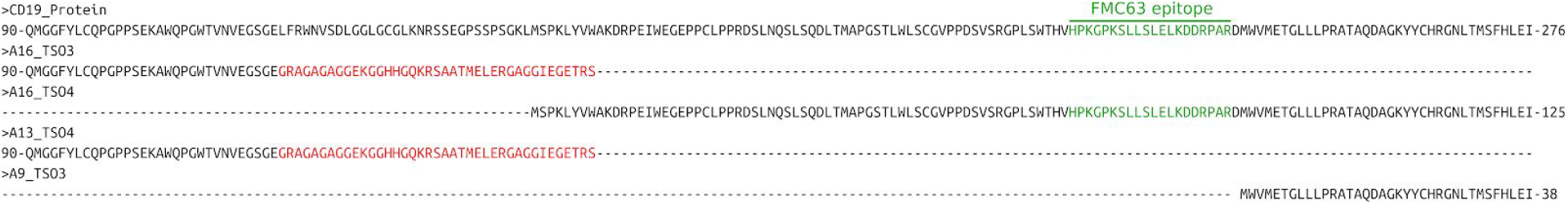
Predicted amino acid sequences of CD19 isoforms lack FMC63 CAR-T cell epitope. CD19 isoforms identified by Mandalorion based on single cell R2C2 consensus reads were manually aligned to the CD19 protein reference (on top). Red letters indicate mismatches to the CD19 protein reference. The FMC63 epitope is shown in green.

### cDNA Generation

**Table S1.**
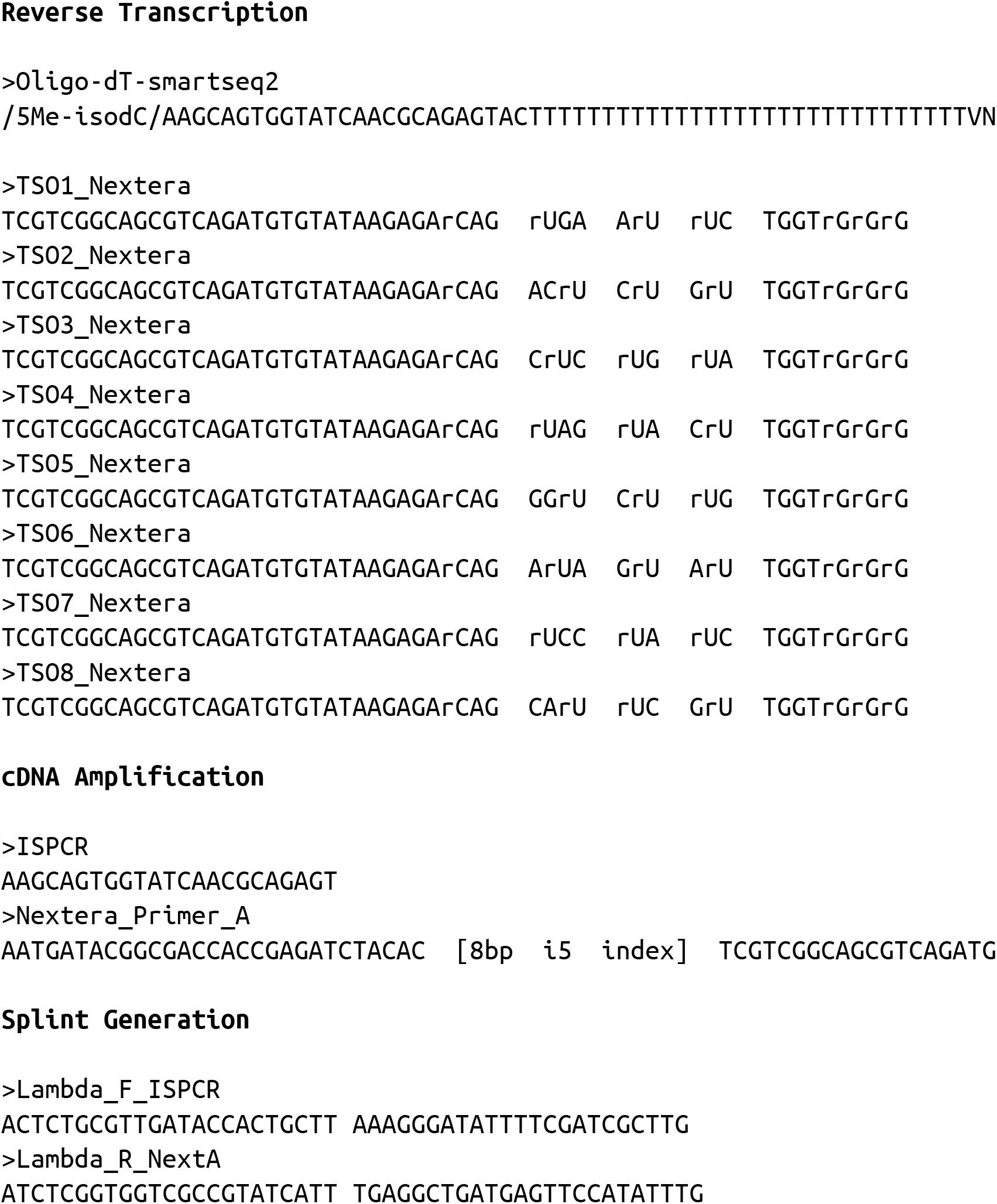
Oligos used in the manuscript. All oligos are shown 5’->3’ and were ordered from Integrated DNA Technologies (IDT). Lower case ‘r’ indicates RNA bases. Spaces in sequences are for visual emphasis only.

